# Neural integrator and orchestrator communities shape spontaneous signaling in the human brain

**DOI:** 10.64898/2026.02.04.703687

**Authors:** Lorenzo Pini, Ranieri Dugo, Paolo Pigato, Maurizio Corbetta

## Abstract

Understanding how intrinsic brain dynamics are organized is critical for explaining human cognition and sensory processing. Theoretical frameworks propose a hierarchical architecture in which some neural systems act as orchestrators, broadcasting information across the brain, whereas others serve as integrators, transforming incoming signals. Here, we quantitatively test whether this orchestration-integration framework is embedded in intrinsic brain activity, that is, in the absence of explicit cognitive/sensorial tasks.

We adopt a multivariate fractional modeling framework originally developed in financial mathematics to characterize how volatility propagates across interacting markets, thereby identifying systems that act as orchestrators or integrators. We then test whether this integrator-orchestrator axis is related to human intelligence. To this end, using 7T resting-state functional magnetic resonance imaging data from 173 healthy young adults, we model spontaneous brain fluctuations with a multivariate fractional Ornstein-Uhlenbeck process to derive directional influence indices.

Consistent with our predictions, we identified a bipartite organization, stable across modeling choices. At the subcortical level, the anterior thalamus, putamen, and caudate emerged as orchestrating transmitters, whereas the posterior thalamus, globus pallidus, hippocampus, amygdala, and nucleus accumbens acted as integrating receivers. At the cortical level, attentional and sensory networks functioned as orchestrating transmitters, while higher-order cognitive networks served as integrating receivers. These findings provide support for a theoretically grounded integration-orchestration framework, demonstrating that brain signaling is organized along this axis even at rest, relevant for intelligence scores. The proposed fractional framework offers a principled tool to investigate how disruptions of this balance may contribute to brain disorders.

**Competing Interest Statement:** None.

## Introduction

The human brain is organized along a hierarchy of anatomical substrates and functional systems.^1–7^ A key challenge in neuroscience is understanding how this hierarchical architecture supports human behavior by governing the directionality of information flow, distinguishing regions that broadcast signals from those that primarily integrate them.

A dominant view is that large-scale coordination of brain activity is carried out by a core subset of brain regions. The classical model by Norman and Shallice^8^ posits that the prefrontal cortex exerts supervisory control over lower-level sensorimotor processes. Similarly, Baars’ global workspace theory^9^ proposes that information is momentarily integrated within a small set of hub regions before being broadcast across the brain. Extending this model, the global neuronal workspace framework by Dehaene and Changeux^10^ suggests that associative areas for perception, memory, attention, and valuation form a high-level unified space where information is not only integrated but also redistributed to lower-level systems. These frameworks all point to a functional distinction between integration, where information is combined, and orchestration, where that information is distributed and influences other regions.

While these models are effective in explaining how the brain operates during active sensory or cognitive engagement, it remains unclear whether a similar hierarchical organization is also expressed during rest, when the brain is not engaged in an explicit task. Addressing this question requires an important conceptual distinction between task-evoked and resting-state neural signals. Task-based paradigms measure fluctuations in neural activity, whether electrical signals recorded with electroencephalography or magnetoencephalography, or hemodynamic proxies such as the blood-oxygen-level-dependent (BOLD) signal measured with functional MRI (fMRI), while participants actively perform a specific task. In these contexts, subtraction paradigms (e.g., task versus baseline or task versus alternative tasks) can be used to identify brain regions or networks (and their functional properties) selectively recruited to support task execution.

In contrast, resting-state paradigms focus on the analysis of neural activity when the brain is not engaged in any explicit task, a condition referred to as “resting-state”. Notably, the resting brain exhibits structured and reproducible patterns of spontaneous activity. A growing resting-state fMRI (rsfMRI) literature has demonstrated that, during rest, the brain maintains coherent activity within specific neural networks, reflecting a form of intrinsic dynamics that closely resembles patterns observed during task performance.^11^ Notably, spontaneous signal fluctuations in sensory cortices recapitulate task-evoked activity patterns. In visual cortex, resting-state fluctuations mirror those elicited during visual tasks engaging the same regions, such as face-selective areas during facial discrimination tasks.^12^ Similar findings have been reported in motor cortex, where spontaneous activity during rest more closely resembles fluctuations associated with familiar, ecologically relevant motor actions than with less naturalistic motor tasks.^13–15^ These observations support the existence of signal reactivations mechanisms during rest, whereby behaviorally relevant neural patterns are reinstated even in the absence of overt task demands.

A fundamental question that follows is whether the theoretical distinction between integrative and orchestrating modules, formulated to explain brain function during task, is also preserved during rest, and whether inter-individual variability in this intrinsic hierarchical organization relates to individual differences in intelligence. In this study we specifically tested the hypothesis that, in the absence of explicit sensory or cognitive demands, the brain retains a directional and hierarchical organization in which certain regions predominantly integrate incoming signals, while others act as orchestrators by broadcasting influence across large-scale networks. Motivated by evidence of replay-like mechanisms during rest,^12–16^ we assessed whether resting-state activity preserves a directional organization compatible with an orchestrator-integrator framework.

We assessed the properties of brain integration and segregation using rsfMRI, a tool capturing brain fluctuations in the absence of task engagement.^17^ Traditionally, these fluctuations are analyzed using static, linear, and univariate models, which, while informative, fall short of capturing the multivariate dynamics observed in brain activity, thus offering a partial view of the mechanisms governing information flow across the brain’s functional architecture.^18–20^

To overcome these limitations, we applied a multivariate fractional model originally developed in financial mathematics to model cross-component dependencies and directional volatility spillovers between market indices.^21^ In the original application to financial systems, the model revealed that some stock indices generally acted as net transmitters of volatility, exerting more influence on other markets than they received. Conversely, the remaining indices consistently functioned as net receivers, responding more to external fluctuations than inducing them.

Drawing a parallel with brain networks, we hypothesize that intrinsic neural activity follows a similarly asymmetric organization, akin to that observed in financial markets. In this framework, fluctuations originating from some brain regions exert a directional influence on the fluctuations of other regions, thereby acting as orchestrators, whereas other regions primarily absorb, retain, and integrate incoming fluctuations, functioning as integrators. In financial systems, such asymmetries emerge naturally from the interaction structure of the market, where certain indices consistently transmit volatility across the system while others predominantly reflect and accumulate these influences. We hypothesized that analogous principles govern intrinsic brain dynamics, enabling the identification of networks that preferentially broadcast neural influence versus those that primarily integrate it.

In principle, alternative multivariate approaches could be used to test this hypothesis.^18^ However, commonly adopted models for dynamic functional connectivity (e.g., dynamic conditional correlation or exponentially weighted moving average frameworks) do not explicitly account for the fractional nature of neural signals, which is increasingly recognized as a defining property of brain dynamics across species.^22–26^ In contrast, the multivariate fractional framework explicitly incorporates temporal dependencies through the estimation of the Hurst coefficient. Specifically, by fitting the Hurst exponent independently for each participant, this approach take into account neural variability, the inherent fluctuations in neural activity arising from multiple intrinsic sources, including stochastic ion channel dynamics, variability in synaptic transmission, and differences in large-scale network processing.^27^ Rather than being mere “noise,” neural variability is shaped by the brain’s structural and functional architecture and manifests across all levels of neural function, from perception to motor output.^28^ Subject-specific estimation of the Hurst coefficient therefore enables the characterization of individual temporal “fingerprints” of neural dynamics, capturing inter-individual differences in signal-memory and persistence within resting-state activity. Consequently, each individual is characterized by a personalized set of model parameters, rather than being constrained to a common group-level solution.

Based on these premises, by treating large-scale brain networks as interacting components of a complex system analogous to financial markets, this multivariate fractional framework was applied to test whether resting-state brain dynamics are organized along an orchestrator-integrator axis, and whether this intrinsic axis is directly related to cognitive performance. Specifically, we hypothesized that this axis would be associated with fluid rather than crystallized intelligence, as the former depends on flexible neural interactions, whereas the latter, reflecting consolidated knowledge, is less reliant on intrinsic neural dynamics. The whole analytical workflow is illustrated in **Fig. 1**.

**Figure 1.**
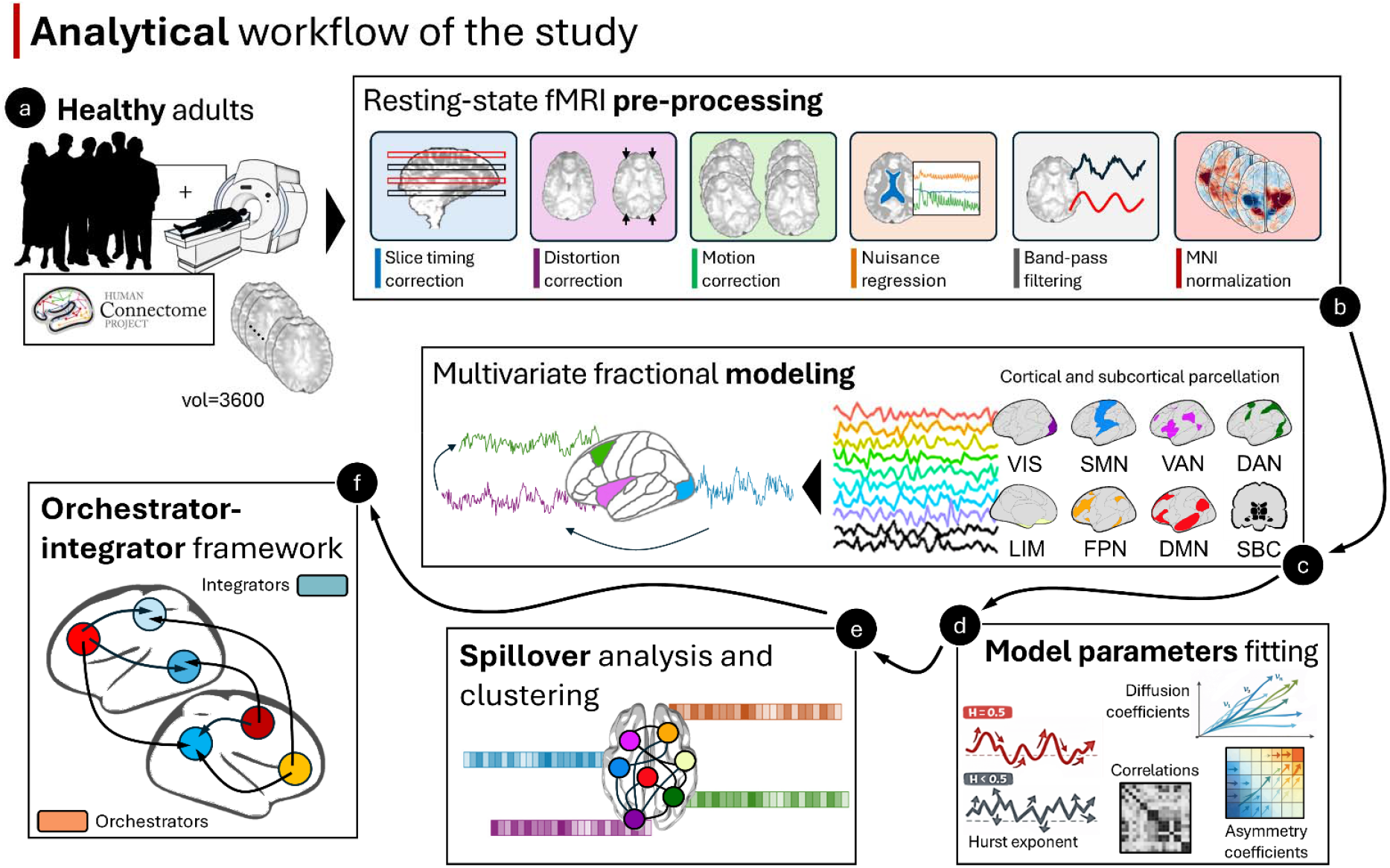
Analytical workflow of the analysis. Resting-state fMRI data from healthy adults of the Human Connectome Project were retrieved (panel a) and preprocessed (panel b). Data were grouped into cortical and subcortical regions by averaging signals according to predefined networks and parcels. A multivariate fractional model was fitted at the individual level to characterize key parameters, including the Hurst coefficient, diffusion coefficients, and correlation and asymmetry coefficients. These parameters were entered into a spillover analysis to identify orchestrator and integrator brain modules using a clustering-based approach.

## Results

In this analysis we employed a large dataset of n=173 healthy adults from the human connectome project (HCP) collected at 7T during rsfMRI. This dataset represents state-of-the-art reference for the assessment of intrinsic brain functional connectivity. The human brain was parcellated according to different regions: the Tian’s parcellation (hippocampus, globus pallidum, anterior and posterior thalamus, caudate, nucleus accumbens, amygdala, and putamen),^29^ and the 7 Yeo’s cortical networks (visual: VIS; sensorimotor: SMN; dorsal-attention: DAN; ventral-attention: VAN; limbic: LIM; default mode: DMN; frontoparietal: FPN).^30^

First, we analyzed the properties of the rsfMRI signal through statistical tests designed to assess stationarity and fractality, deeming appropriate the modeling of the BOLD fluctuations with the multivariate fractional Ornstein-Uhlenbeck (mfOU) process, able to reproduce these two behaviors, further allowing for a rich interdependence structure, and the fitting of the time-series on an individual basis.

### The model parameters

The N-dimensional mfOU process models a system in which each component (here network and subcortical parcels) is the result of two forces: a driving Gaussian noise, which is fractional and correlated across components, and a mean-reverting dynamic that pushes the process towards a certain mean value. More formally, we have that the Gaussian noise is characterized by four parameters: the vector of Hurst coefficients (H), *H* ∈ [0,1]^*N*^ which regulates the fractality (or complexity); the vector of diffusion coefficients, *v* ∈ *R*^*N*^,which determines the level of variability and coincides with the square root of variance at time 1; the matrix of correlations, *ρ* ∈ [−1,1]^*N*×*N*^, where each element *ρ*_*i,j*_ determines the contemporaneous correlation between components i and j; and the matrix of asymmetry coefficients, *η*, which determines the asymmetry of the lagged (non-contemporaneous) correlations. *H*_*i*_ ∈ [0, 1], *i* = 1, …, *d*, deliver a Markovian process when 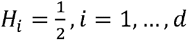, rough marginal trajectories (high complexity) when 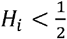 and smoother ones (lower complexity) when 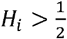. On top of the noise, the mfOU process entails mean-reverting dynamics ruled by the vectors of the speed of mean reversion coeficients *α* ∈ *R*^*N*^, and the long-term means *μ* ∈ *R*^*N*^. We assume *μ* = 0. In addition, to study spillovers among cortical and subcortical brain regions, we assumed causality, i.e. that the current values depend on the past Gaussian noise, which prescribes

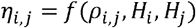

for a certain function *f*, which is why this parameter won’t be presented among the estimates (*see section below*).

### Brain macro-region estimates

Before employing the model, we evaluated the stationarity of the signal using the Dickey-Fuller test and its fractionality by means of the Change of Frequency estimator.^31^ When testing the null hypothesis of non-stationarity, we get p-values<0.01 for all regions across all individuals, providing strong evidence in favor of the mean-reverting nature of the time series. The COF estimates are shown in **Supporting Fig. S1**, which highlights the tendency of the brain signals towards roughness (high complexity), as H fall below 0.5. The Tian’s regions (subcortical regions and the hippocampus, hereafter called subcortical for simplicity) appear clearly rough (H<0.5). Only for one individual in the hippocampus and twelve individuals in the putamen H was not significantly smaller than 0.5. At the cortical level H took values at or above 0.5 on most nodes, except for the LIM, which sits on the rough side of the spectrum. A notable degree of inter-subject variability was evident (**Supporting Fig. S1**), underscoring the appropriateness of the individual fits to capture potential sources of neural variability in the rsfMRI signal, relevant for the patterns under investigation.

**Supporting Fig. S2** shows the parameters of the mfOU process estimated on the Tian’s regions across all individuals by Minimum Distance Estimation (MDE), which identifies the model parameters by minimizing the distance between model and empirical covariances. The speed of mean reversion parameters is estimated below 0.5 in most individuals and, overall, around 0.26. Given *δ* = 1(rsfMRI time repetition unit), these values correspond to apparent mean-reverting behaviors over the time span we consider. Anterior thalamus, putamen, and caudate show faster mean reversion (*α*_*i*_ ≥ 0.35) than the other regions (∼0.2). The diffusion coefficients, which are estimated to be around 40, show the local variability of the signals, and tell that some regions such as nucleus accumbens (∼50) display more variability in the BOLD signal than others, such as hippocampus (∼36). The values in *v*_i_ are quite consistent across left-right hemisphere pairs. H, which still points towards roughness, differs from the COF estimates shown in **Supporting Fig. S1** due to the different estimation strategy and are now higher. This difference is because the COF estimator only considers the regularity of the time-series while the MDE estimator fits its covariance. **Fig. 2** represents the matrix *ρ* as a heatmap. From this map it is possible to appreciate that the two hemispheres of each node are characterized by the high values, with the exception of globus pallidus (0.11). Low values are observed for inter-node pairs that include hippocampus, amygdala and globus pallidus. Overall, the group made of anterior thalamus, putamen, and caudate presents the highest values of the speed of mean reversion coefficients, low roughness (high regularity), and high *ρ*_*i,j*_ s. On the other hand, the globus pallidus is characterized by lower speeds of mean reversion, lower H, and lower contemporaneous dependencies from the driving noise.

**Figure 2.**
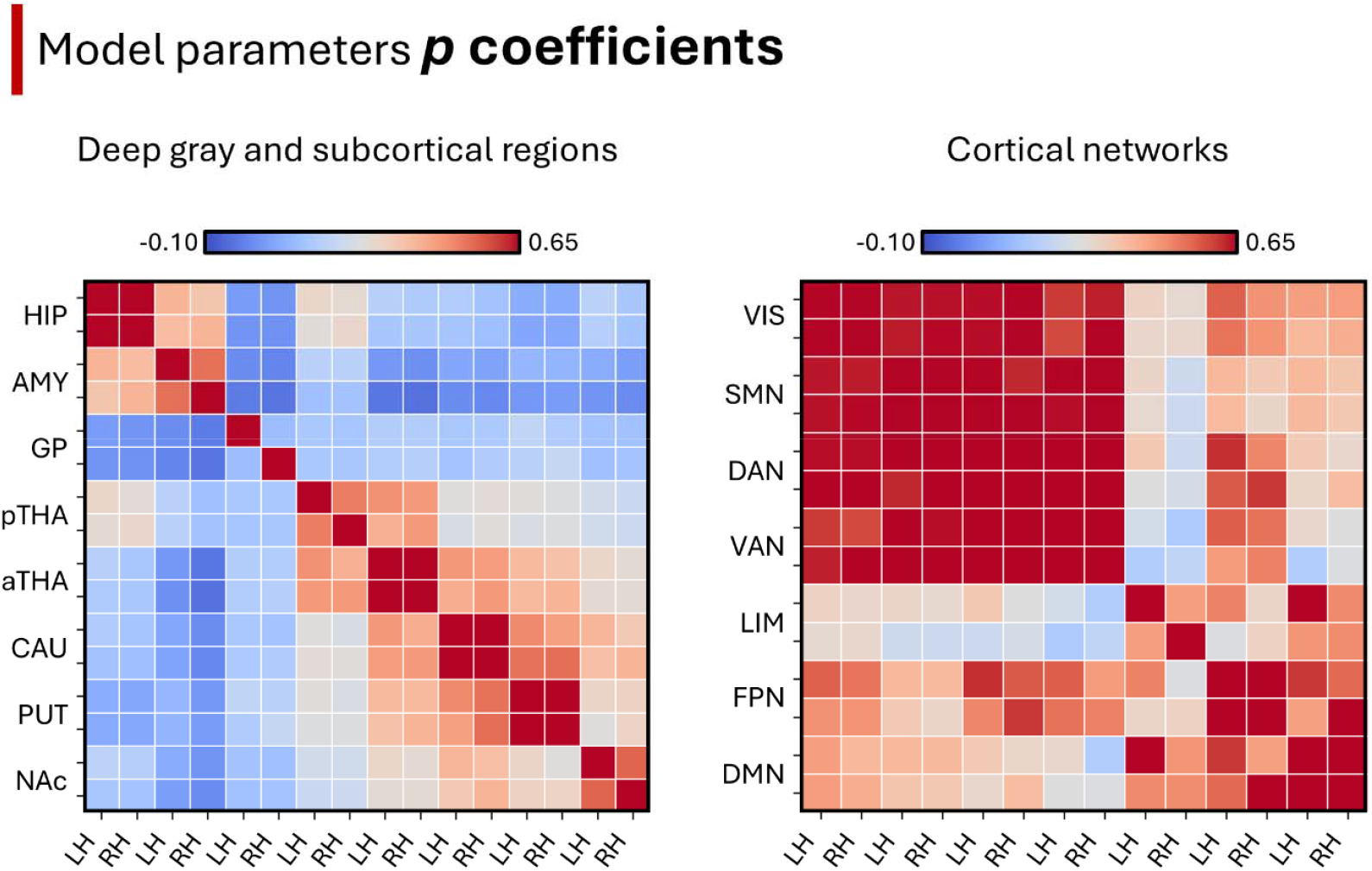
Deep gray, subcortical structures and cortical network communities. Left panel: p parameter matrix across subcortical regions and deep gray regions. Right panel: p parameter matrix across cortical networks. Abbreviations: HIP: hippocampus; AMY: amygdala; GP: globus pallidus; pTHA: posterior thalamus; aTHA: anterior thalamus; CAU: caudate; PUT: putamen; NAc: nucleus accumbens: VIS: visual network: SMN: sensorimotor network; DAN: dorsal-attention network: VAN: ventral-attention network; LIM: limbic network; FPN: frontoparietal network: DMN: default mode network; LH: left hemisphere; RH: right hemisphere. The upper bound of the color scale was set to 0.65 to improve contrast between regions; higher values (e.g., diagonal) are saturated.

Similar parameters of the mfOU process are reported for the aggregated cortical networks (**Supporting Fig. S3**). The estimates of the univariate marginal parameters in **Supporting Fig. S3** show two key facts: (i) speeds of mean reversion take values like their subcortical counterparts, except for the higher values and variabilities in the LIM node, which drive them a bit higher overall (∼0.3), and (ii) markovianity is not ruled out. Regarding the latter, estimates of the Hurst coefficients are now more centered around 0.5, suggesting that a simple model driven by a standard Brownian motion might be a good fit for these time-series. Nevertheless, the H estimates for LIM are quite spread, making it difficult to reject its roughness. The t-tests could not reject the hypothesis H=0.5 for all cortical nodes. However, the H estimates for LIM deviate from normality, making it difficult to assess the markovianity of the signal. Therefore, we provided a comparison of spillovers with a simpler Markovian model to assess the robustness of the results. Similarly to the subcortical regions, the cortical networks also demonstrate notable inter-individual variability, highlighting the importance of accounting for subject-specific differences in the modeling approach (**Supporting Fig. S3**). The values of *v*_*i*_ were lower, mostly around 30, with exception for LIM (∼14).

Regarding the values in *ρ* (**Fig. 2**), we observed high values between different hemispheres of the same region, more than in the subcortical parcels (mostly above 0.7). Qualitatively, VAN, DAN, SMN, and VIS form a cluster of high *ρ*s, while DMN, FPN, LIM seem to form another one driven by less correlated noise. This impression was confirmed by the spillover analysis discussed in the following paragraph (*see below*).

### Subcortical and Cortical Spillovers

Spillovers are a method for quantifying lead-lag relationships in time-series exploiting the concept of the forecast error variance decomposition, which was first introduced in Econometrics.^32^ The forecast error variance decomposition determines how much variability in forecasting a component of the system, i.e. calculating its future expectation, is attributable to the innovations in another component of the system occurring during the forecast horizon. This idea encapsulates the delayed influence of the dynamics of one component over another and corresponds to the building block denoted directional pairwise spillover. Aggregating the pairwise spillovers appropriately, is it possible to recover the influence exerted among groups of variables. For example, the total influence received by one component from all the others (received spillovers), the total influence that a component exerts on all the others (transmitted spillovers), and the corresponding net quantities corresponding to the difference between transmitted and received ones. Definitions are given in Section *Spill-over analysis*. We apply this framework to the mfOU process and its estimated parameters on subcortical and cortical brain signals.

Due to the lack of normality of spillover values we performed a Friedman X^2^ test with pairwise non-parametric post-hoc analyses (Bonferroni corrected) for comparing net spillovers (defined as the difference between transmitted and received spillovers for each region).

At the subcortical level, the analysis revealed a significant region effect across individuals (left hemisphere: F=688; p<0.0001; right hemisphere: F=798; p<0.0001)(**Fig. 3**). Results echoed for both the left and right hemispheres. For simplicity the post-hoc analysis was performed on the left-right average values. We found significant region differences, with the globus pallidus and the anterior thalamus at the lowest and highest extreme, respectively (p<0.001 Bonferroni corrected). Only hippocampus compared to the posterior thalamus (p=0.175), and amygdala compared to the nucleus accumbens (p=0.120) showed no significant results (**Fig. 3**).

**Figure 3.**
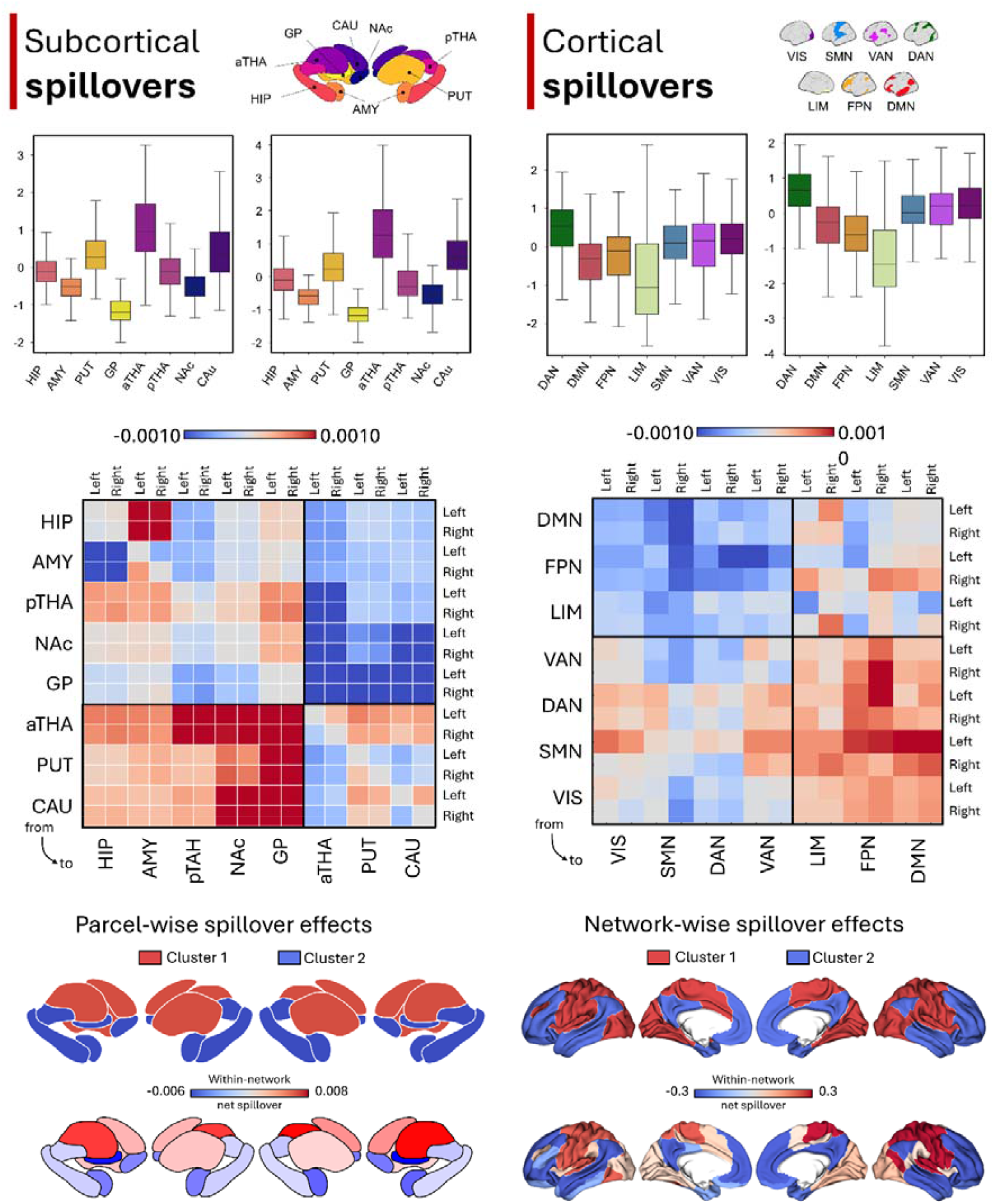
Net Spillover patterns. Top panels: Net spillover in subcortical regions (including the hippocampus) across the entire sample (left); net spillover within Yeo’s 7 cortical networks across the entire sample (right). Median and standard deviation are reported. Each color represents a brain region, as illustrated in the legends above the panels. Center panels: net spillover effects between subcortical (left) and cortical networks (right). Blue: negative net effects; red: positive net effects. Bottom panels: net effects projected at subcortical (left) and cortical (right) surface level; the top panels depict the clustering between network (red: cluster 1; blue: cluster 2); bottom panels depict the column-wise matrix average net values for each cortical network.

At the cortical network level, we found a significant network effect for both the left (F=179; p<0.0001) and right hemispheres (F=293; p<0.0001)(**Fig. 3**). Post-hoc analysis run considering the left-right average net-spillover values showed significant differences among most of the comparisons (p<0.001 Bonferroni corrected). Non-significant differences were found between VIS and SMN (p_uncorrected_ = 0.062), VIS and VAN (p_uncorrected_ = 0.123), SMN and VAN (p_uncorrected_ = 0.548), LIM and FPN (p_uncorrected_ = 0.141), and between DMN and FPN (p_uncorrected_ = 0.583). A significant difference not-surviving multiple corrections was found between LIM and DMN (p_uncorrected_ = 0.004).

### Spillovers communities

We performed a clustering analysis of the net spillover pattern at the aggregated-level (considering average-spillover values across the whole cohort of individuals) to identify the brain communities grouped into receivers (integrators) and transmitters (orchestrator) through a k-means analysis. The net spillover effect defined two main clusters (defined by the consensus clustering). At subcortical level, an ‘orchestrator cluster’ was composed of the anterior thalamus, putamen, and caudate and a ‘integrator’ one by hippocampus, amygdala, posterior thalamus, nucleus accumbens and globus pallidus. Results are shown on **Fig. 3**.

At cortical-network level we observed two main clusters, as shown by the consensus clustering analysis. The “orchestrator” community was composed by DAN, VAN, SMN, and VIS. On the other hand, the “integrator” community was characterized by FPN, DMN, and LIM (**Fig. 3**).

Results were largely replicated using hierarchical factor analysis (HFA), conducted to assess the robustness of the identified communities. In the Tian subcortical parcellation, the first factor (42% variance explained) loaded primarily on the anterior thalamus, putamen, and caudate, bilaterally. The second factor (39% variance explained) loaded on the hippocampus, posterior thalamus, nucleus accumbens, and globus pallidus, bilaterally. The bilateral amygdala exhibited low loadings on both factors. Similarly, for the network parcellation, HFA showed two main factors, the first loading mainly on bilateral VIS, SMN, DAN, and VAN (47% variance explained), and the second mainly on left LIM, and bilateral FPN, and DMN (33% variance explained). Results are presented in **Fig. 4**.

**Figure 4.**
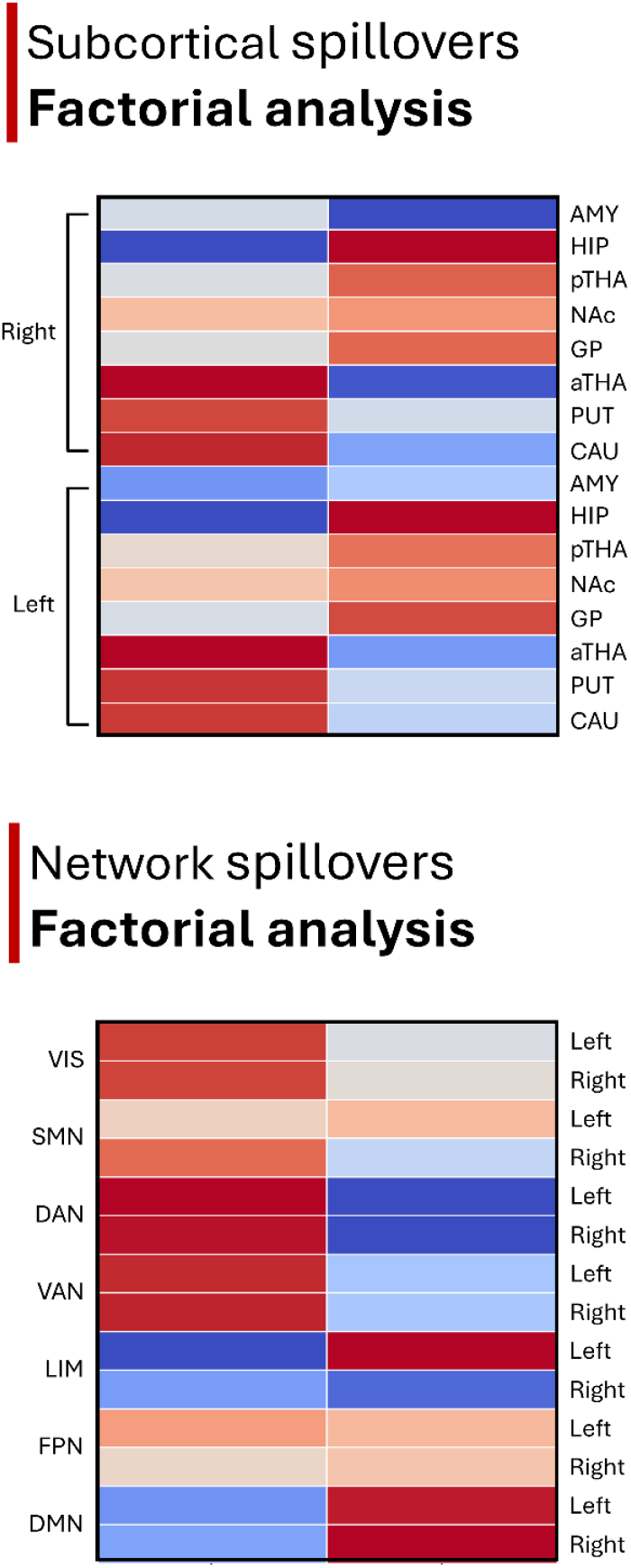
Factorial analysis on net Spillover. Loading values for the first two main factors from net spillover values for subcortical parcellation (top panel) and cortical networks (bottom panel).

### Brain intrinsic Integration-orchestration axis and human intelligence

To assess the behavioral relevance of the integration-orchestration framework during resting-state, we investigated the relationship between net spillover scores grouped according to the orchestrator and integrator communities (computed as median effect from both cortical network and subcortical parcels) with cognitive performance, focusing on two measures of intelligence: fluid and crystallized. Fluid intelligence score was positively associated with integration (r=0.21, p=0.007), although the effect was modest, whereas orchestration showed no significant correlation (r =-0.09, p=0.23). No associations were observed for crystallized intelligence scores (integration: r=0.04, p=0.58; orchestration r=0.07, p=0.37). Results are shown on **Fig. 5**. Linear models confirmed that for fluid intelligence, integration showed a significantly stronger effect than orchestration (t=2.70, p=0.008), indicating that variation in integration more strongly predicts fluid intelligence. For crystallized intelligence, no significant difference was observed between integration and orchestration (t=0.88, p=0.38).

**Figure 5.**
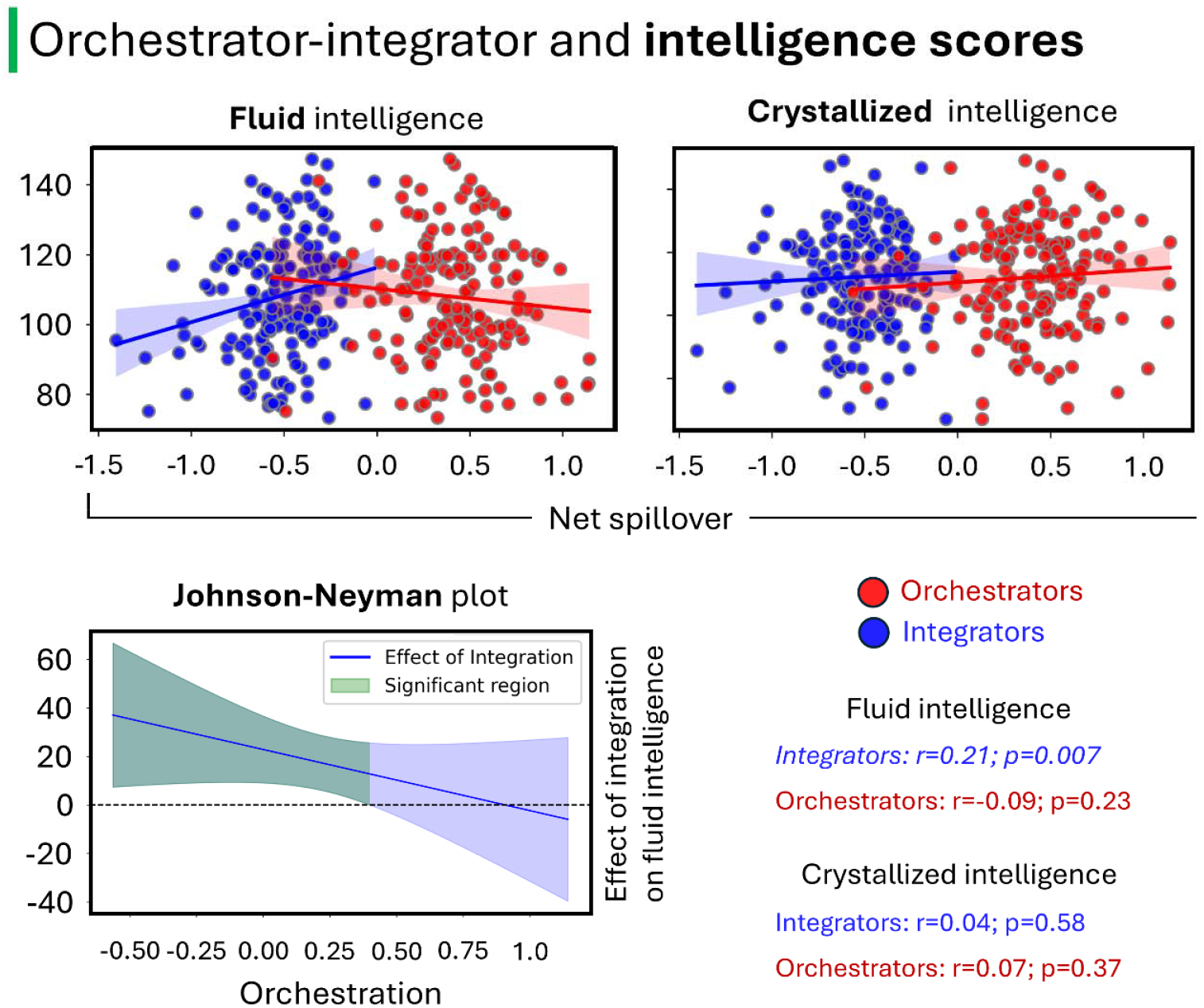
Relation between brain intrinsic orchestration-integration axis with fluid intelligence. Top panels: simple relation between net spillover in brain communities clustered as integrators (blue) or orchestrators (red) for both fluid (left) and crystallized (right) intelligence performance. Bottom panel: Johnson-Neyman plot for the simple slope of integration predicting fluid intelligence. The relation was significant when orchestration was lower but not significant with higher scores of orchestration.

The Johnson–Neyman analysis revealed that the effect of integration on fluid intelligence was significant only at low levels of orchestration. At higher levels of orchestration, the relationship attenuated and was no longer significant (**Fig. 5**), indicating that integration contributes to fluid cognition when orchestration is not maximal.

### Robustness of the cortical network spillover across the sample

To evaluate the robustness of the net spillover results across individuals, we conducted a sensitivity analysis aimed at quantifying the consistency of the observed patterns at the individual level, limited to the cortical network structure. Using the group-average spillover pattern as a reference, both the Frobenius distance and the Structural Similarity Index (SSIM) yielded very low (consistent) values across the sample (Frobenius: mean=0.009: min=0.006; max=0.014; SSIM: mean=. 982; min=0.96; max=0.99), indicating high overall similarity. Among the networks, the LIM system exhibited the greatest inter-individual variability, although the absolute distance values remained relatively low. A k-means clustering analysis based on both Frobenius and SSIM values identified n=4 individuals forming a distinct cluster, characterized by consistently higher distance values. These individuals were interpreted as outliers and represent less than 3% of the total sample, thereby supporting the high stability and reproducibility of the results across participants (**Fig. 6**).

**Figure 6.**
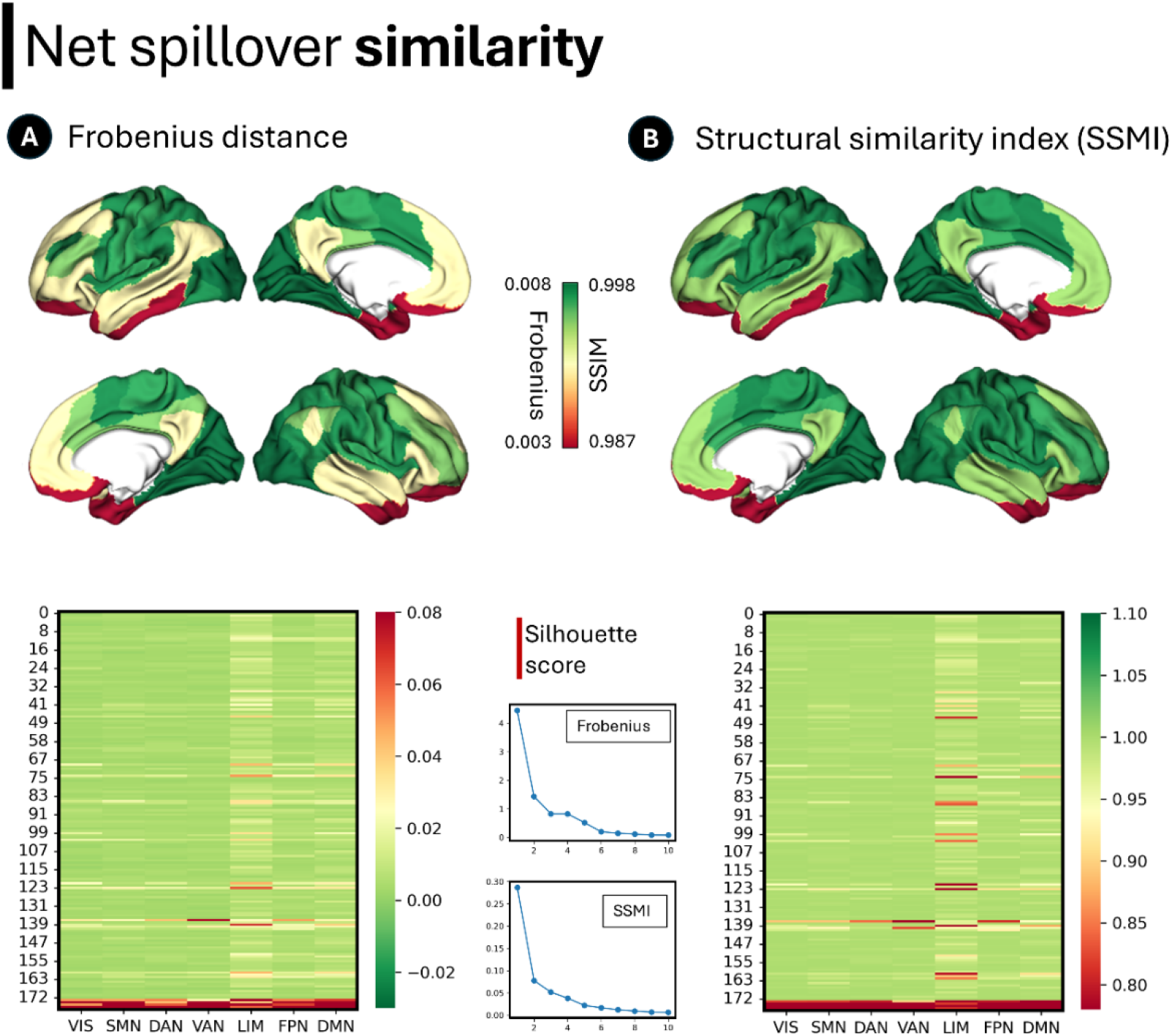
Net spillover similarity across individuals. For each individual, the similarity to the group-average net spillover pattern was quantified using the Frobenius distance (panel A) and the Structural Similarity Index (SSIM; panel B). The top panels display the averaged similarity measures projected onto the brain surface, where green indicates higher inter-individual similarity and red indicates lower similarity. The bottom panels show network-wise distributions of Frobenius distance and SSIM values for each individual. Data are presented ordered according to the two groups identified by the clustering analysis using the similarity indexes.

### Model choice and robustness of results

The results described are based on the multivariate, fractional model recently introduced in Dugo et al.^21^ This modelling framework is justified by i) the fractal properties of the measured signal,^19^ ii) the presence of signal-memory, and iii) the scale-free dynamics in brain activity.^33,34^ In principle, lead-lag relationships could be quantified with a spillover analysis based on different model specifications. For example, the standard framework,^35,36^ is based on Vector Autoregressive (VAR) modelling. To get a measure of robustness of our results, we performed a similar analysis using VAR in place of mfOU. Notably, we find that the two frameworks mostly agree for cortical networks and partially agree on subcortical ones. Regarding the former (**Supporting Fig. S4**), both methodologies identify DAN and LIM at the two extremes of the orchestrator-integrator axis. They mainly agree also with the results on the other networks. The two methodologies also display similar magnitudes. Regarding subcortical networks (**Supporting Fig. S5**), the VAR framework echoed the results on anterior thalamus as a prominent orchestrator and the globus pallidus a prominent integrator, although higher magnitudes were found for the putamen as orchestrator and posterior thalamus and nucleus accumbens as integrator. The directionalities were consistent except for hippocampus. Overall, VAR results showed similar results reporting similar clusters within the orchestrator-integrator framework. To be note, since the signal shows fractional properties, according to a recent analysis a fractional model is considered more appropriate for the aim of this study,^37^ thus the subcortical results should be interpreted in the framework of mfOU.

## Discussion

Our findings offer empirical evidence in support of the integration-orchestration framework during resting-state.^38^ Previous theories, such as the global neuronal workspace and supervisory attentional systems,^9,10^ posit a functional division between high-level integrative hubs and lower-level processing systems. Here, we advanced this view by providing a directionally explicit map of how neural influence is distributed across brain cortical and subcortical regions at rest. By applying spillover analysis based on describing the rsfMRI signals with the mfOU process, we identified specific brain modules.

At the subcortical level, we observed a community suggestive of “orchestrators” modules, comprising the anterior thalamus, caudate, and putamen; on the other hand, the globus pallidus, hippocampus, amygdala, nucleus accumbens, and posterior thalamus might function as “integrators.” These results are consistent with classical models of cortico–basal ganglia–thalamocortical loops. The caudate and putamen, as primary input nuclei of the basal ganglia, are known to relay cortical signals toward the globus pallidus and thalamus.^39,40^ Notably, the analysis reported a different role for the anterior and posterior thalamus, with the former showing the highest transmitter profile among subcortical/deep gray regions. These findings may challenge the view of the thalamus as a universal information receiver,^41^ instead suggesting a differentiated role for its posterior and anterior portions during resting-state that operate in concert to facilitate information integration.^42^ The amygdala and hippocampus, while traditionally linked to limbic and memory systems, are increasingly recognized for their integration within these circuits through the ventral striatum and thalamic pathways.^43–45^ Our spillover analysis offers a functional perspective that aligns with this directional architecture, highlighting the anterior structures as putative drivers and posterior-limbic nuclei as integrative receivers. Interestingly, when examining the spillover patterns of the hippocampus and amygdala in isolation (see **Fig 4.a**), the former appears to act as a orchestrator. Although the clustering analysis places these two structures within the same module, this potential dynamic aligns with recent evidence suggesting that some insular sites and the hippocampus can exert reciprocal influence, either modulate memory processes or providing inputs that become encoded as episodic memory traces. Clarifying the functional significance of this hippocampus–amygdala directional relationship, and its potential interplay with insular pathways, represents a compelling avenue for future research.^46^

At the cortical network level, we described a broader dichotomy, encompassing two principal communities of networks. Specifically, the DAN, VAN, VIS, and SMN formed a cluster of “orchestrators,” while the DMN, FPN, and LIM clustered as “integrators.” This bipartite organization aligns with the theoretical framework of cortical processing gradients,^2^ in which unimodal and attentional systems preferentially convey information (transmitter net spillovers), whereas transmodal networks integrate and contextualize these inputs within internal models, capturing the functional asymmetry in information flow. These results are in line with a recent study applying energy-landscape analysis to examine the resting-state brain dynamics. While this study did not examine the causal interplay between networks, two main states were observed in healthy controls. A first state dominated by activation of LIM, FPN, and DMN, while DAN, VAN, VIS, and SMN were inactive, and a second state in which active-inactivation patterns were reversed.^47^ These states in patients with degenerative conditions were altered, suggesting that the dynamics of these brain states could be a critical biological endophenotype. The large-scale functional segregation likely reflects, and is constrained by, the underlying cytoarchitecture: primary sensory regions exhibit dense granular layers, supporting input-driven processing, whereas association cortices tend to be agranular, reflecting their role in integrating information and generating abstract representations.^48^ These gradients underpin a spectrum of functional specialization, whereby lower-level regions primarily receive and process external stimuli, and higher-level areas generate predictions and internal models.

From a neurobiological standpoint, SMN and VIS are tightly coupled to external sensory and motor functions and are known for their rapid, stimulus-driven dynamics.^49^ VAN and DAN, while part of higher-order attentional systems, are also closely linked to detecting salient stimuli and reorienting attention, respectively. Their role as orchestrators may reflect their function in dynamically broadcasting task-relevant or salience-tagged information across the cortex.^50^ While the VAN has been associated with bottom-up reorienting and salience detection, and the DAN supports sustained top-down attentional control, the joint categorization we found in this study could reflect the shared property of rapidly disseminate context-relevant signals to drive perception and behavior across the cortex replayed during resting-state conditions.

In contrast, DMN, FPN, and LIM are transmodal and integrative in nature. The DMN has been extensively implicated in internally directed cognition, autobiographical memory, and self-referential processing; the FPN in executive control and working memory; and the LIM system in emotion, interoception, and semantic knowledge. Our results suggest that these systems operate by accumulating, integrating, and recontextualizing signals originating from sensory-attentional networks. In particular, the DMN and LIM have been hypothesized to contribute to the “internal workspace” of consciousness, integrating external stimuli with stored representations and internal bodily states.

Overall, this core of orchestrating and integrative brain regions aligns with previous theories proposing a global neural workspace in which sensory, motor, attentional, and memory systems interconnect to form a higher-level unified space.^51^ However, our findings clearly distinguish attentional–sensory networks from higher-order cognitive networks (e.g., those involved in memory), thereby refining this framework at rest. This pattern is consistent with results from Deco et al. (2021), who used transfer entropy applied to task-fMRI data to demonstrate that outgoing information flow (drivers) arises primarily from unimodal sensory regions, whereas incoming flow (integrative receivers) is concentrated in higher-order transmodal and limbic regions.^38^

Beyond large-scale network organization, the integration–orchestration axis showed a selective relationship with human behavior. Higher intrinsic integration was associated with better fluid intelligence. Crucially, the effect of integration depended on orchestration levels: integration positively predicted fluid intelligence only when orchestration was relatively low, with the relationship attenuating at higher orchestration. This pattern suggests that fluid cognition benefits from efficient integration of distributed information, but only within a balanced regime of large-scale signal broadcasting. The absence of any association with crystallized intelligence further supports the specificity of this intrinsic axis for flexible, adaptive cognitive functions rather than accumulated knowledge.

Importantly, the clustering of networks into transmitter and receiver modules also parallels observations in pathological conditions. Recently, we have identified DAN as the core network in the context of consciousness disorders, namely Anton syndrome and anosognosia for hemiplegia.^52^ Patients exhibiting visual or motor deficits along with anosognosia (i.e., a lack of awareness of their impairments) showed overlapping patterns of structural disconnection involving the DAN, when compared to patients with similar deficits who retained full awareness.^52^ Overall, these observations support the hypothesis that the DAN operates not only as a mediator of sensory-guided attention but also as a key orchestrator of signal propagation across the cortex. From a systems-level perspective, the DAN may serve as a dynamic relay that conditions the functional state of downstream networks by broadcasting top-down signals, in line with global neuronal workspace theories.^51,53^

Conversely, dysfunctions in the DMN and LIM systems have been linked to impairments in self-awareness, interoception, and affective regulation, as seen in depression, and disorders of consciousness.^54^ This suggests that the directional dynamics observed in the healthy brain are not only theoretically meaningful but may also carry clinical relevance as markers of network integrity and functional hierarchy.

Unlike traditional approaches that infer hierarchy from static connectivity or task engagement, our mfOU-based spillover analysis allowed us to move beyond general descriptors of integration and to quantify the asymmetry of neural variability, revealing that specific systems consistently act within an orchestration-integration axis even at rest.

These results were stable using a VAR methodology. A recent analysis in Bibinger et al.^37^ points out that VAR models may not perform well when applied to fractional time series that involve persistent (or anti-persistent) memory. Further, in the VAR framework, the spillovers depend on the lag of the autoregressive process as well as a forecast horizon to be fixed. For this reasons we opt for our model,^21^ as spillover indices do not depend on such parameters.

Despite the robustness of the proposed framework, several limitations should be acknowledged. First, the analysis relied exclusively on 7T rsfMRI data from healthy young adults in the HCP, which may limit the generalizability of the findings to other groups or lower-field acquisitions. Second, the temporal resolution and intrinsic hemodynamic properties of fMRI constrain the direct inference of neuronal activity, potentially introducing biases in the estimation of memory-properties and spillover indices. Finally, the parcellation schemes adopted, though widely validated, impose *a priori* spatial boundaries that may obscure finer-grained heterogeneity.

Overall, this work offers a unifying perspective on brain hierarchy, setting the stage for future studies examining how this directional architecture is altered in clinical populations or across development.

## Materials and Methods

### Sample and fMRI preprocessing

In this study, we included n=173 healthy individuals from the HCP (female 61%; age range: 22-36; range 22-25 years, n=20; range 26-30 years, n=84; range 31-35 years, n=67; +36 years, n=2). Participants underwent a 7T MRI examination. Data were pre-processed according to the minimal pre-processing HCP pipeline. Briefly, the data were slice-timing corrected, whole-brain intensity normalized, distortion corrected using synthetic field map estimation and spatially realigned within and across rsfMRI runs. Data were regressed to remove sources of spurious variance from head motion (six parameters), the average signal over the whole brain, ventricles, and CSF, and white matter. These masks were identified through Structural MRI segmentation using Freesurfer (<u>surfer.nmr.mgh.harvard.edu</u>). Finally, temporal filtering was applied to retain frequencies in the 0.009–0.08-Hz. Frame censoring was computed using framewise displacement with a threshold of 0.5 mm and images were normalized to the MNI space through a non-linear approach.

### Testing for stationarity and fractionality

In order to motivate our modeling choice, we performed formal stationarity and fractionality tests. The Augmented Dickey-Fuller test tests the null hypothesis that a unit root is present in a time series sample. The alternative hypothesis is stationarity.

When looking at fractionality, we relied on the change of frequency (COF) estimator considered in Barndorff-Nielsen et al.^31^ The COF estimator is semiparametric and relies only on the assumptions of stationarity and Gaussianity. It also comes with a sound asymptotic theory that delivers associated standard errors.

Due to the nature of our timeseries, we consider it appropriate to model the rsfMRI signal with the mfOU process. It is worth noting that the estimates’ densities for H presented in **Supporting Fig. S1** differ from those in **Supporting Fig. S2 and Supporting Fig. S3**. This is due to the fact that the former are obtained with the COF estimator, which looks at the geometrical characteristics of the series using their numerical derivatives, whereas the latter are obtained with the MDE estimator, which relies on the law of the process by targeting cross-covariances.

### The model

The model we used in the study was recently introduced mathematically and applied to volatility timeseries data in our previous publications.^21,55^ It is a multivariate mean-reverting fractional process, that we refer to as the multivariate fractional Ornstein-Uhlenbeck (mfOU) process. Specifically, it represents a stationary version of the multivariate fractional Brownian motion (mfBm) introduced by Amblard et al.^56,57^

Let 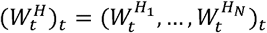 be a mfBm, i.e. a vector-valued Gaussian process in dimension *N* with parameter *H* ∈ (0,1)^*N*^, *ρ* ∈ ⌈−1,1⌉^*N*×*N*^ and *η* ∈ ℝ^*N*×*N*^. In this context, *H* = (*H*_1_, …, *H*_*N*_) is the vector of Hurst exponents of the components, the matrix *ρ* represents the contemporaneous correlation of the mfBm at each point in tim and the time matrix *η* rules the asymmetry in time of the cross-covariance function and the time reversibility of the process. These matrices are such that *ρ*_*i,i*_ = 1, *ρ*_*i,j*_ = *ρ*_*j,i*_, *η*_*i,i*_ = 0, and *η*_*i,j*_ = − *η*_*j,I*_ for *i,j* …, *N*.

The mfOU process 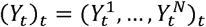,is the vector process whose components are the stationary solutions to equations

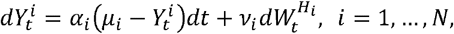

where *μ*_*i*_ ∈ ℝ is the long-term mean, *v*_*i*_ > 0 is the diffusion coefficient, and *α*_*i*_ > 0 is the speed of mean reversion, i.e., larger *α*_*i*_ produce dynamics that stay closer to the long-term-mean, and are in a sense more stable, while smaller *α*_*i*_ correspond to a more dispersed dynamics, and *α*_*i*_ = 0 characterizes a non-stationary process (mfBm). This process is Gaussian, meaning that it is completely defined by its covariance matrix and as a byproduct of fractionality, its value at every time step is correlated to its infinite past. A more detailed mathematical description is given in Dugo et al.^58^ We measure time in TR, in accordance with the sampling frequency of the data.

### Estimation

We estimate the parameters of the model using a MDE.^59^ We assume to have *n* equally spaced observations of the process over the interval [0,T], that we use to compute sample auto and cross-covariances with time lags 0,1,2,3,4,5,10, and 15 TRs. Then, we chose as parameter the ones minimizing the distance between sample and model-implied cross-covariances, in a mean-square sense. This estimator is consistent and asymptotically normal, with speed of convergence √*n* if *max* (*H*_1_,…,*H*_*N*_) < 3/4. The estimator and its properties are described in detail in our previous publication.

### Spill-over analysis

Spillovers are defined in a multivariate system of time series as the shares of the forecast error variance in one variable due to the effect of the innovations in another variable. Let *ψ* _*i,j*_ (*h*), be the share of the variance in the error of predicting 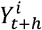 due to innovations in the variable 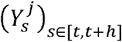, for *i,j* = 1,…, *N*. In Pesaran (1997)^60^, in the discrete time setting, this is defined as

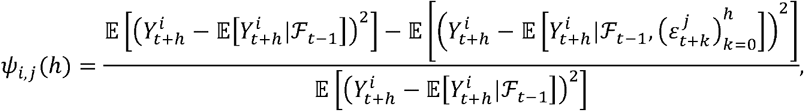

where *ε*_*t*_ is the *N*-variate white noise innovation in *Y*_*t*_ at time 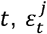 is its *j*-th component, and ℱ_*t*-1_ is the information set at time *t* − 1, i.e. the filtration generated by the underlying *ε*_*t*−*i*_,*i* = 1,…, To use the information in the variance decomposition matrix Ψ, with entries [Ψ(*h*)]_*i,j*_ = *ψ*_*i,j*_ (*h*), in order to construct spillover indices in the presence of correlated white noises, consider the normalized quantities

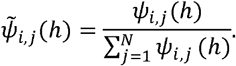

In Dugo et al.^21^ we provided the following representation for the quantities above, for the adapted time discretization of the causal mfOU process, assuming t>h:

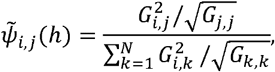

where

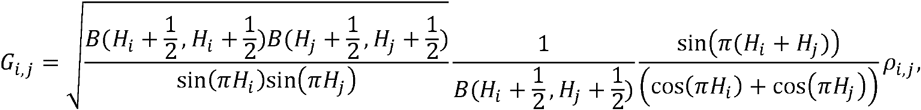

*and B*(*x,y*), *x,y* > 0, *denotes the Beta function*. The 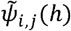 are aggregated according to the rules outlined in **Supporting Table S1** to construct the spillover indices.

### Net spillover analysis

Due to the non-normality of the spillover distribution (Shapiro-Wilk test^61^), we applied the Friedman test (non-parametric alternative to repeated-measures ANOVA)^62^ to assess whether significant differences existed across brain regions in terms of net spillovers. To investigate pairwise differences, post-hoc Wilcoxon signed-rank tests were performed for all region pairs.^63^ To control multiple comparisons, p-values were corrected using the Bonferroni method. This procedure was applied independently for both the subcortical and cortical structures.

Further, we analyzed the clustering structure of the net spillover group-average matrix. First, to identify the optimal number of clusters we applied a consensus clustering procedure.^64^ For each candidate number of clusters (from k=2 to 7), we performed 100 independent k-means runs with a single initialization to account for variability in clustering solutions. For each run, a co-association matrix was computed, indicating the proportion of times each pair of samples was assigned to the same cluster. These matrices were averaged across runs to form a consensus co-association matrix for each k. Hierarchical clustering (average linkage) was then applied to the distance matrix derived from the co-association matrix, and the resulting clusters were evaluated for stability using the Silhouette score (from scikit-learn library: https://scikit-learn.org/stable/index.html). The number of clusters with the highest stability was selected as optimal. Final cluster assignment was obtained by running k-means with the optimal k.

To assess the robustness of the clusters, we run an additional HFA to the same average net spillover matrix. Cortical measures were first standardized using z-scores. HFA was performed on the standardized data extracting two factors using the principal axis method and a Varimax rotation, following our previous publications.^65,66^ Eigenvalues and factor loadings were then obtained to assess the contribution of each cortical measure to the extracted factors.

### Integration-orchestration with intelligence

Composite indices of network organization were computed by taking the median across relevant regions based on integrators and orchestrators communities (see section *Net spillover analysis*). Cortical and subcortical measures were then combined to generate overall integration and orchestration indices by averaging the corresponding cortical and subcortical components. Extreme outliers in these composite indices were removed using an interquartile range criterion, excluding values beyond 1.5 times the IQR from the 5th and 95th percentiles. Linear relationships between the network median-composite measures and intelligence performance (fluid and crystallized composite scores) were assessed using Pearson correlation coefficients. Linear regression analyses were conducted to examine the independent and interactive effects of overall integration and orchestration on intelligence performance, separately for fluid and crystallized scores using ordinary least squares regression. Interaction terms between integration and orchestration were included to assess whether their combined influence differed from their individual effects.

To further explore the interaction between overall integration and orchestration on intelligence, a Johnson-Neyman analysis was conducted.^67^ Specifically, we tested the hypothesis that the effect of integration on fluid intelligence performance varies across different levels of orchestration. To this end, simple slopes of integration were calculated across the observed range of orchestration values. Standard errors and 95% confidence intervals for these slopes were computed.

### Sensitivity analysis (network-level)

Finally, we quantified the similarity between the net spillover pattern with the reference group-average matrix. For each subject, we computed both Frobenius norm and structural similarity index (SSIM) between the individual net-spillover matrix and the reference (group-average). This procedure allows to compute distance values for each individual in each network. The resulting matrix (m×n; with n representing individuals and m representing networks) was input into a clustering algorithm to identify potential subgroups of participants. A higher dispersion of individuals across different clusters would indicate lower consistency within the sample. This index was used as measure of reliability of the retrieved brain communities.

## Acknowledgments

Data were provided by the Human Connectome Project, WU-Minn Consortium (Principal Investigators: David Van Essen and Kamil Ugurbil; 1U54MH091657) funded by the 16 NIH Institutes and Centers that support the NIH Blueprint for Neuroscience Research; and by the McDonnell Center for Systems Neuroscience at Washington University.

## Author Contributions

LP: conceptualization, discussion, data analysis, interpretation of results, writing; RD: data analysis, discussion, writing; PP: supervision, conceptualization, discussion, proofreading; MC: supervision, proofreading, discussion

## Supporting Information

**Fig. S1.**
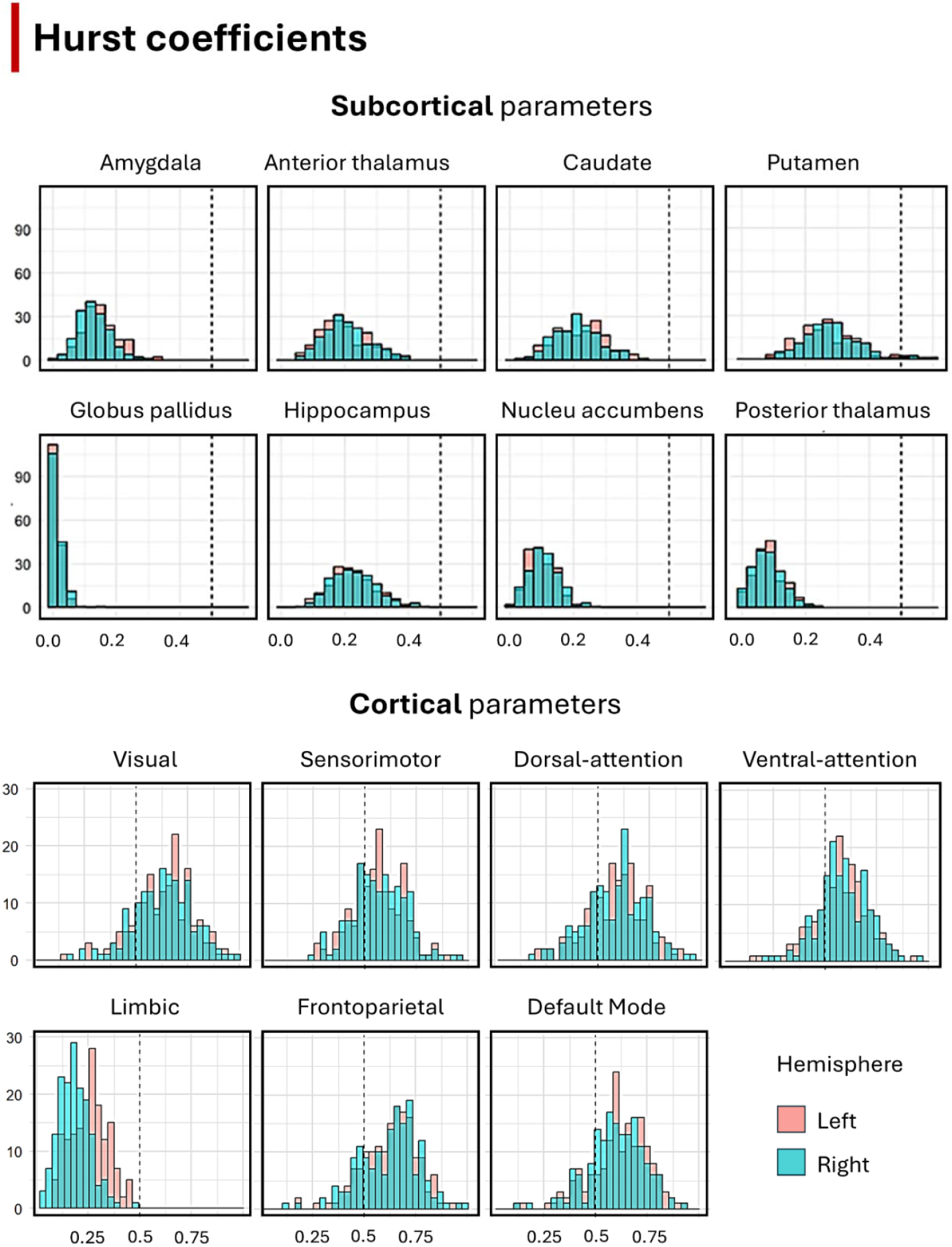
Estimates of the Hurst coefficients obtained with the Change of Frequency (COF) estimator.^1^ Top panel: subcortical regions (plus hippocampus). Bottom panel: cortical regions.

**Fig. S2.**
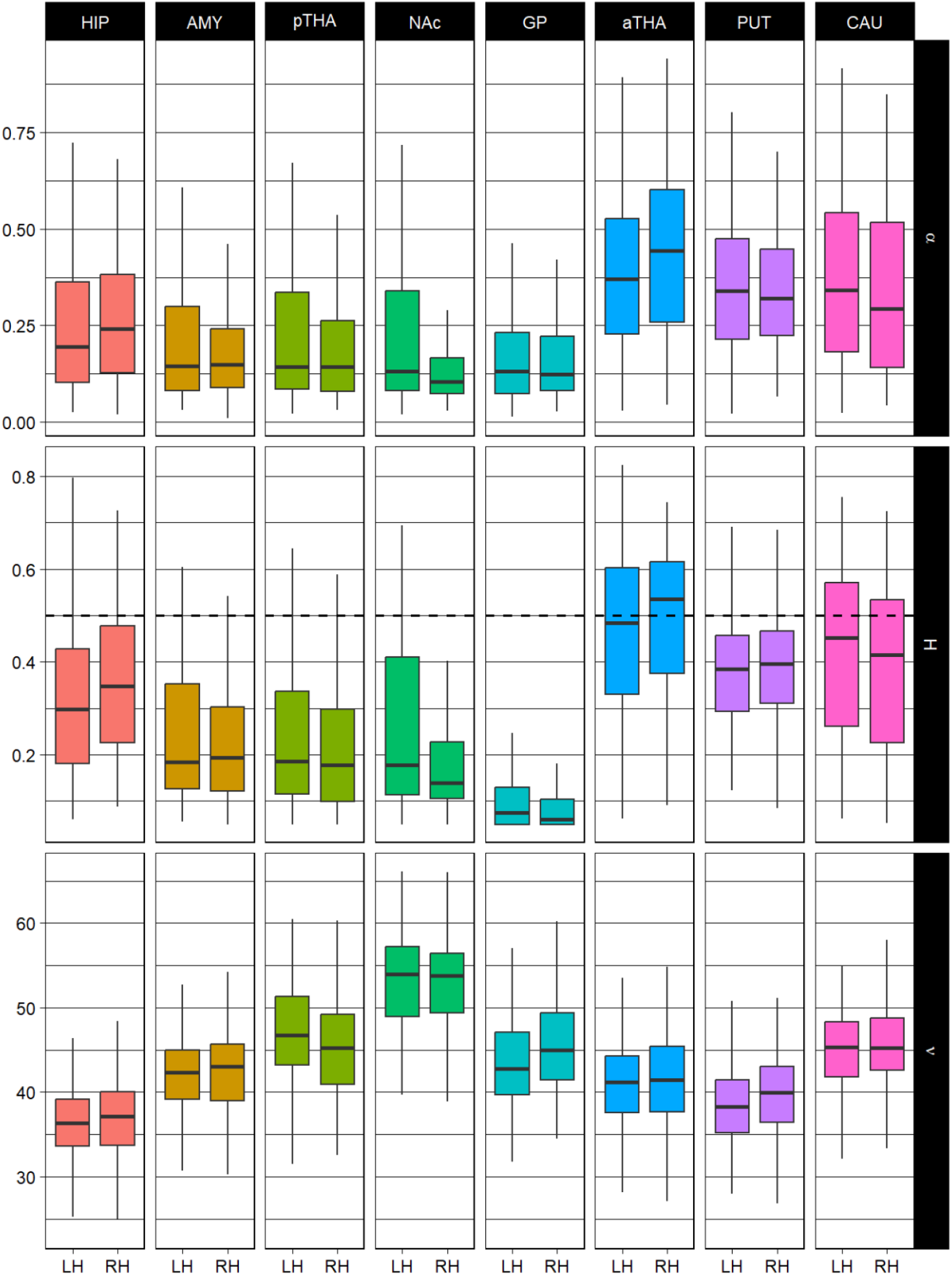
Boxplot representation of the parameter estimates of the mfOU process applied to the resting-state functional MRI signals of individuals’ subcortical and hippocampal parcels. Abbreviations: HIP: hippocampus; AMY: amygdala; pTHA: posterior thalamus; NAc: nucleus accumbens; GP: globus pallidus; aTHA: anterior thalamus; PUT: putamen; CAU: caudate; LH: left hemisphere; RH: right hemisphere.

**Fig. S3.**
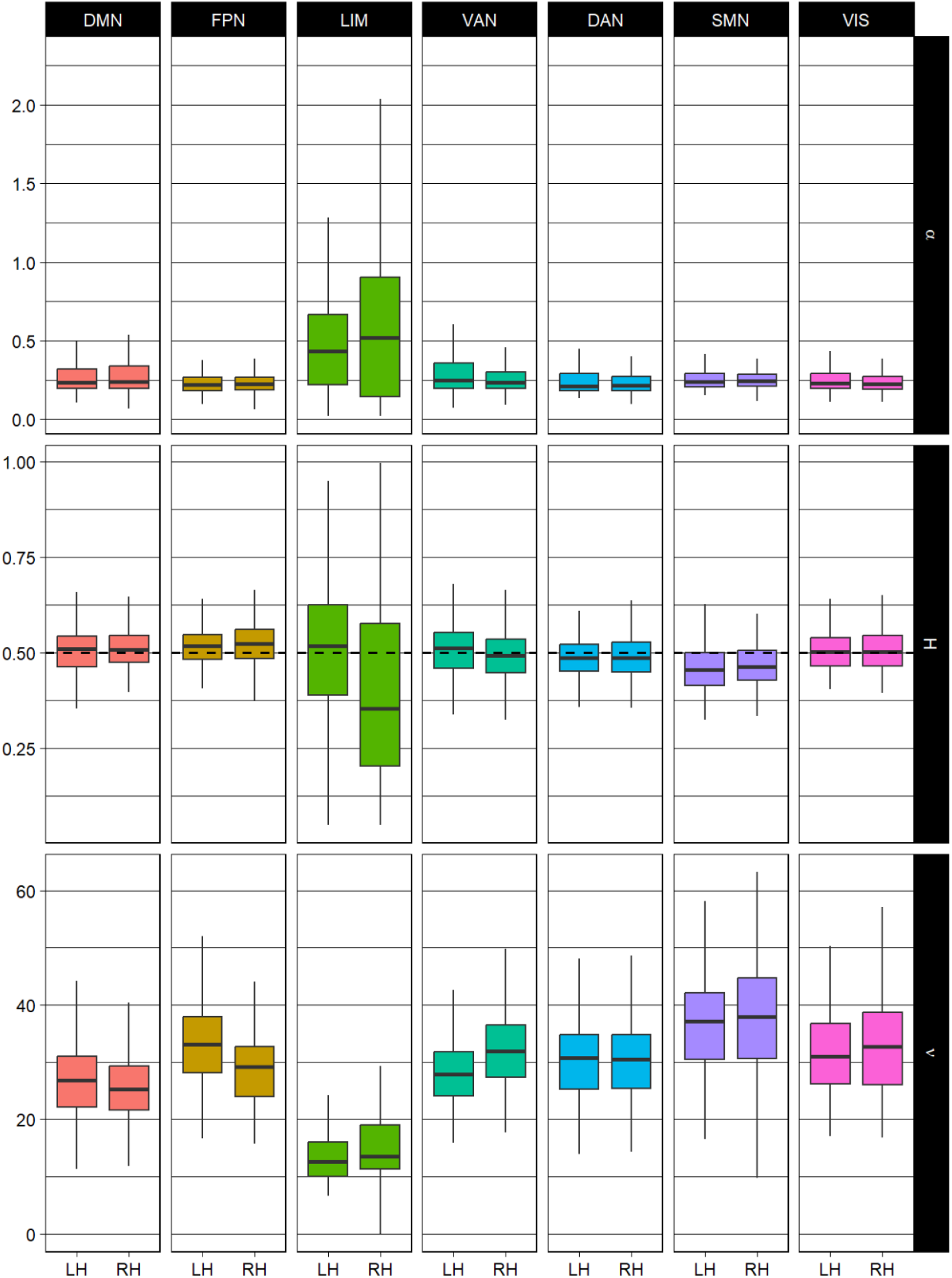
Boxplot representation of the parameter estimates of the mfOU process applied to the BOLD signals of individuals’ cortical networks (aggregate measures). Abbreviations: DAN: dorsal-attention network; DMN: default-mode network; FPN: frontoparietal network; LIM: limbic network; SMN: sensorimotor network; VAN: ventral-attention network; VIS: visual network; LH: left hemisphere; RH: right hemisphere.

**Fig. S4.**
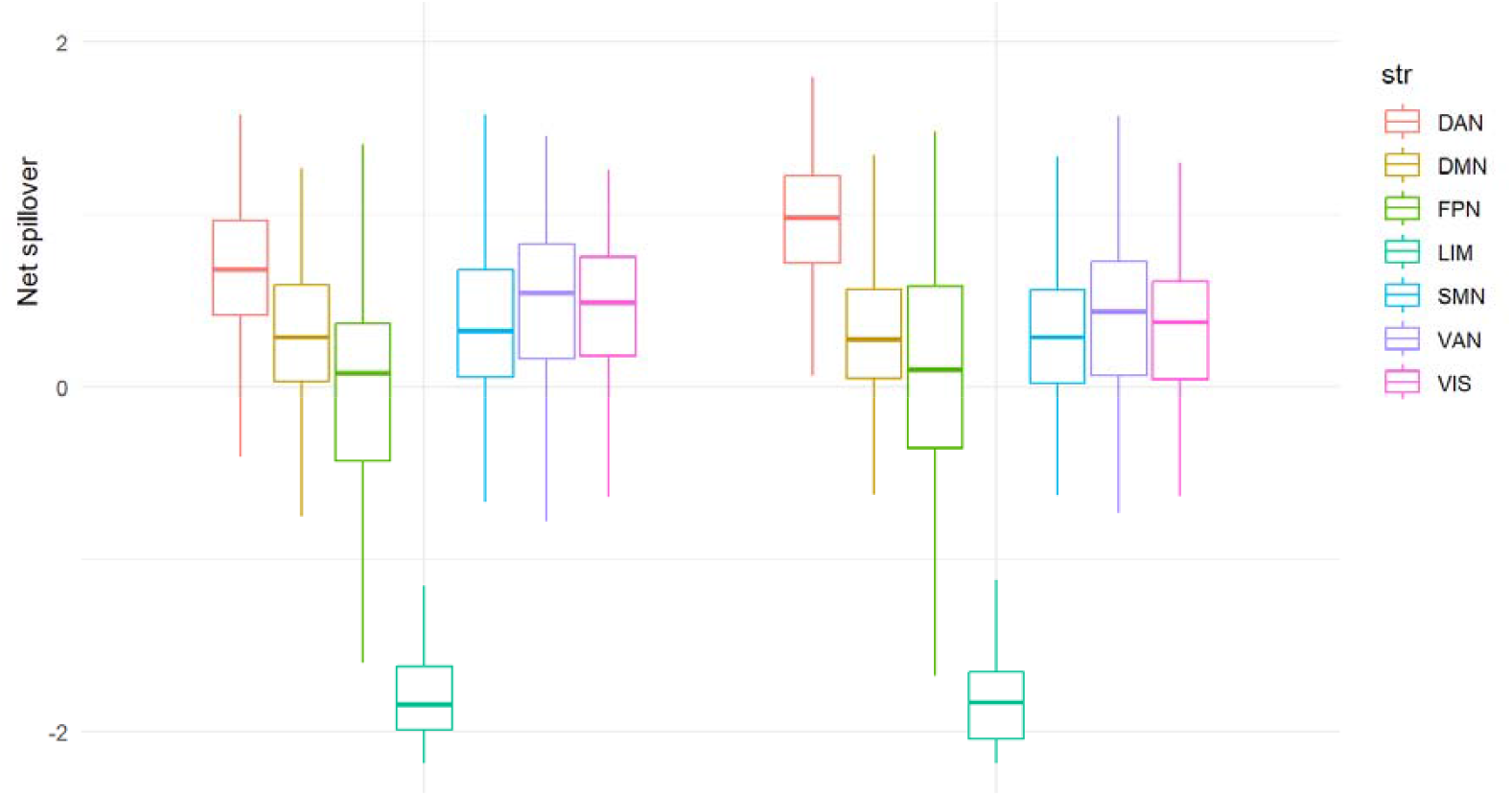
Boxplot representation of the net spillover measures based on VAR modeling of the BOLD signals of individuals’ cortical networks (aggregate measures). Left plots represent the left hemisphere; right plots represent the right hemisphere. Abbreviations: DAN: dorsal-attention network; DMN: default-mode network; FPN: frontoparietal network; LIM: limbic network; SMN: sensorimotor network; VAN: ventral-attention network; VIS: visual network.

**Fig. S5.**
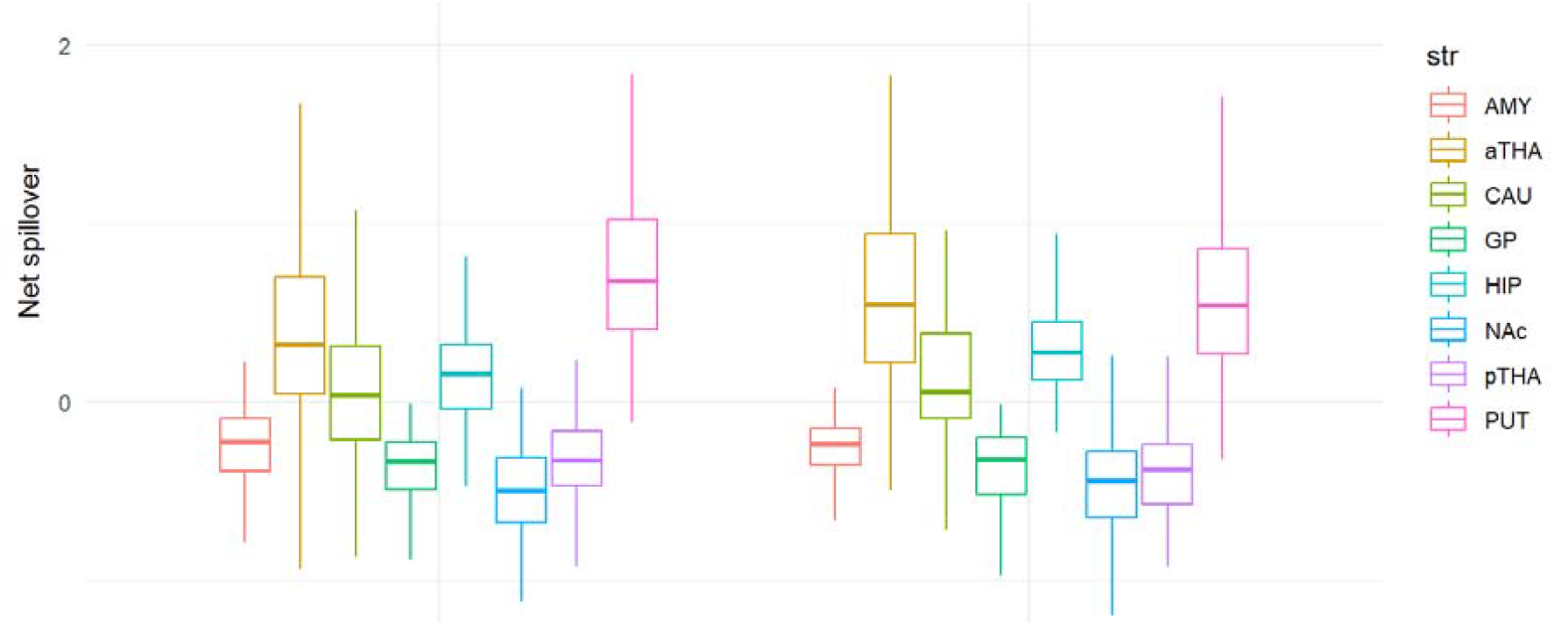
Boxplot representation of the net spillover measures based on VAR modeling of the resting-state functional MRI signals of individuals’ subcortical and hippocampal parcellation. Left plots represent the left hemisphere; right plots represent the right hemisphere. Abbreviations: HIP: hippocampus; AMY: amygdala; pTHA: posterior thalamus; NAc: nucleus accumbens; GP: globus pallidus; aTHA: anterior thalamus; PUT: putamen; CAU: caudate.

**Table S1.** Definitions of Spillover Indices from Diebold and Yilmaz^2^, where 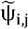 (h) is defined in Section 4.5 of the main text.

| Spillover Index | Description | Definition |
| --- | --- | --- |
| Total | Aggregation over all components in the system of the normalized forecast error variances due to shocks in other components. | $S(h) = \frac{\sum_{i,j=1, i \neq j}^N \tilde{\psi}_{ij}(h)}{N} \cdot 100$ |
| 1-3 Received | Share of total spillovers received by a component of the system from all the other components. | $S_{i,\cdot}(h) = \frac{\sum_{j=1, j \neq i}^N \tilde{\psi}_{ij}(h)}{N} \cdot 100$ |
| 1-3 Transmitted | Share of total spillovers transmitted by a component of the system to all the other components. | $S_{\cdot,i}(h) = \frac{\sum_{j=1, j \neq i}^N \tilde{\psi}_{ji}(h)}{N} \cdot 100$ |
| 1-3 Net | Difference between the spillovers transmitted those received for each component. | $S_i(h) = S_{\cdot,i}(h) - S_{i,\cdot}(h)$ |
| 1-3 Net Pairwise | Difference between the spillovers transmitted from component $i$ to component $j$ and those transmitted from $j$ to $i$. | $S_{i,j}(h) = \left( \frac{\tilde{\psi}_{ji}(h) - \tilde{\psi}_{ij}(h)}{N} \right) \cdot 100$ |

